# Evolution and variation of 2019-novel coronavirus

**DOI:** 10.1101/2020.01.30.926477

**Authors:** Chenglong Xiong, Lufang Jiang, Yue Chen, Qingwu Jiang

## Abstract

**Background:** The current outbreak caused by novel coronavirus (2019-nCoV) in China has become a worldwide concern. As of 28 January 2020, there were 4631 confirmed cases and 106 deaths, and 11 countries or regions were affected.

**Methods:** We downloaded the genomes of 2019-nCoVs and similar isolates from the Global Initiative on Sharing Avian Influenza Database (GISAID and nucleotide database of the National Center for Biotechnology Information (NCBI). Lasergene 7.0 and MEGA 6.0 softwares were used to calculate genetic distances of the sequences, to construct phylogenetic trees, and to align amino acid sequences. Bayesian coalescent phylogenetic analysis, implemented in the BEAST software package, was used to calculate the molecular clock related characteristics such as the nucleotide substitution rate and the most recent common ancestor (tMRCA) of 2019-nCoVs.

**Results:** An isolate numbered EPI_ISL_403928 showed different phylogenetic trees and genetic distances of the whole length genome, the coding sequences (CDS) of ployprotein (P), spike protein (S), and nucleoprotein (N) from other 2019-nCoVs. There are 22, 4, 2 variations in P, S, and N at the level of amino acid residues. The nucleotide substitution rates from high to low are 1·05 × 10^−2^ (nucleotide substitutions/site/year, with 95% HPD interval being 6.27 × 10^−4^ to 2.72 × 10^−2^) for N, 5.34 × 10^−3^ (5.10 × 10^−4^, 1.28 × 10^−2^) for S, 1.69 × 10^−3^ (3.94 × 10^−4^, 3.60 × 10^−3^) for P, 1.65 × 10^−3^ (4.47 × 10^−4^, 3.24 × 10^−3^) for the whole genome, respectively. At this nucleotide substitution rate, the most recent common ancestor (tMRCA) of 2019-nCoVs appeared about 0.253-0.594 year before the epidemic.

**Conclusion:** Our analysis suggests that at least two different viral strains of 2019-nCoV are involved in this outbreak that might occur a few months earlier before it was officially reported.

## Background

Coronaviruses (CoV) (order Nidovirales, family Coronaviridae) are enveloped, positive stranded RNA viruses, which includes 4 genera *Alphacoronavirus* (α-CoV), *Betacoronavirus* (β-CoV), *Gammacoronavirus* (γ-CoV), and *Deltacoronavirus* (δ-CoV) [1,2]. Since 1960, six different CoVs have been identified and two epidemic CoVs have emerged in human during the last 2 decades [3]. Severe acute respiratory syndrome (SARS)-CoV in 2002-2003 and the Middle East respiratory syndrome (MERS)-CoV in 2012-2015 both belong to the genus *Betacoronavirus* [4,5].

On 31 December 2019, Wuhan City of Hubei Province reported a number of cases of pneumonia related to a local seafood market. The main clinical manifestation of these cases was fever, a few patients had dyspnea, and the chest films showed double-lung infiltrative lesions [6]. The Chinese authorities identified a new type of coronavirus (novel coronavirus, 2019-nCoV), which was isolated on 7 January 2020, and on 11 and 12 January, the World Health Organization (WHO) received further detailed information from the National Health Commission about the outbreak. As of 28 January 2020, there were 4631 confirmed cases and 106 deaths, and had affected 11 countries or regions [7,8].

All human CoVs may be of zoonotic origin, and may indeed originate from bats [9]. To discover the evolutionary history of the viruses, we analyzed the genomes of 2019-nCoVs, compared their variations in both nucleotide and amino acid sequence levels, and determined their molecular clock related characteristics such as the nucleotide substitution rate and the most recent common ancestor. We tried to explore the origin and evolution mechanism of the viruses causing the epidemic, and to provide theoretical basis for early warning of similar events caused by β-CoVs in the future.

## Methods

### Genomes collection

Complete Genomes of 2019-nCoVs were downloaded from the Global Initiative on Sharing Avian Influenza Database (GISAID, http://platform.gisaid.org/epi3/frontend) on January 24, 2020. The SARS-CoV in 2002-2003 (GenBank accession: NC_004718) and two Bat SARS-like coronavirus isolates bat-SL-CoVZC45 (GenBank accession: MG772933) and bat-SL-CoVZXC21 (GenBank accession: MG772934) were also downloaded for serving as the reference sequences. The former is a reference sequence (RefSeq) of pathogens that have caused major epidemics [10], while the latter two are the wild strains with the highest identity with 2019-nCoVs [11]. Moreover, unlike GISAID, the three sequences downloaded from GenBank have detailed comments, which can be used to investigate the coding sequences (CDS) of different proteins.

### Phylogenetic tree construction and amino acid residues alignment

The CLUSTAL W profile alignment option was used to align the whole genomes. On this basis, the coding sequence (CDS) of ployprotein (P, ORF1ab), spike protein (S), nucleoprotein (N), matrix protein (M) and small envelope protein (E) in the genomes are trimmed out from each reference sequence, since they are very important enzymatic or structural proteins in coronavirus [10,12]. Pairwise distances were calculated by using the DNADIST program of the PHYLIP package and by using the Kimura two-parameter model of nucleotide substitution. Phylogenetic trees were constructed by using neighbour-joining (NJ) methods. All phylogenetic trees were inferred by using MEGA6 software [13]. The deduced amino acid residues of these 5 proteins were compared by Lasergene 7 program again [14].

### Computation of mean evolutionary rate and the most recent common ancestor (tMRCA)

Bayesian coalescent phylogenetic analysis, implemented in the BEAST v1.6.1 (http://beast.bio.ed.ac.uk) software package, was used to determine the molecular evolutionary rate [15]. Because the viral genomes sequenced in this outbreak are relatively concentrated in time, no reference sequence other than the outbreak has been included.

GTR+G+I multiple models were chosen as the nucleotide substitution and site heterogeneity models. Each analyses consisting of 10,000,000 generations under the strict clock model because the sequence identity in each nexus matrix is very high. Bayesian skyline population size was set as demographic models for each species-specific data set. Model selection was based on an analysis of marginal likelihoods, calculated in Tracer version 1.5. Effective sample sizes (ESSs) of at least 200 were considered appropriate and the results were considered valid. For tMRCA analysis, the collection dates of 2019-nCoVs were calculated by using the following formula, and the values of date were accepted by Tipdate time collector:

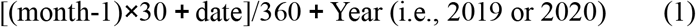

## Results

### Phylogenetic trees and genetic distances

As of January 24, 2020, there were 26 viral genomes of 2019-nCoVs. After excluding one with incomplete information (EPI_ISL_402125) and another excessively short one (EPI_ISL_402126), a total of 24 viral genomes were included in the follow-up study.

The phylogenetic trees showed that all isolates of 2019-nCoVs automatically clustered on whole genome and CDS of five proteins, indicating their close relationship. However, in the whole genome and the CDS of P, S and N, an isolate (EPI_ISL_403928) is different from other 23 (**Fig 1**). This isolate was derived from a 61-year-old male patient in Wuhan. The sample was collected on 1 January, 2020, and the possibility of false sequencing was ruled out. The genetic distances of genomes show results consistent to the phylogenetic trees. The mean genetic distance is 0.00021±0.00002 for all the isolates but only 0.00010±0.00001 after an exclusion of EPI_ISL_403928. The mean genetic distance between EPI_ISL_403928 and the other 23 is 0.00146±0.00018. The CDS of P, S and N show a similar difference between EPI_403928 and the group of 23 (**Tab 1**). All isolates of 2019-nCoV however, have the same CDS of M and E.

**Fig 1.**
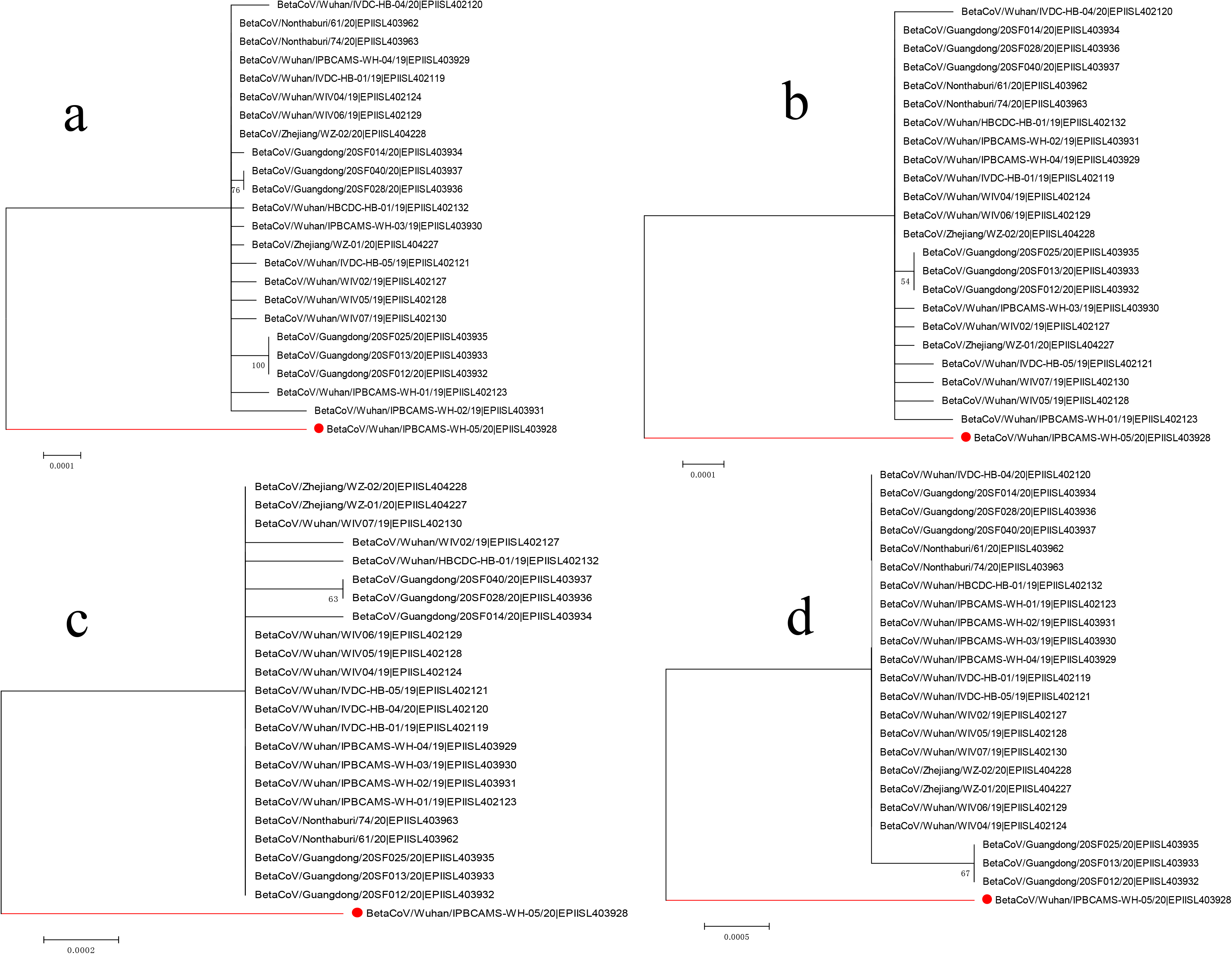
Phylogenetic trees of 24 isolates of 2019-nCoV. a- d correspond to the whole length genome, the coding sequences (CDS) of polyprotein (P), the spike protein (S), and nucleoprotein (N). EPI_ISL_403928 is labelled by red dot and line.

**Tab 1.** Genetic distances of 24 isolates of 2019-nCoV*.

|  | Overall mean d.* | Within mean Gp d. |  | Between Gp mean d. |
| --- | --- | --- | --- | --- |
|  |  | Gp# 1 | Gp 2 |  |
| Genome | 0.00021±0.00002 | 0 | 0.00010±0.00001 | 0.00146±0.00018 |
| P | 0.00018±0.00003 | 0 | 0.00007±0.00002 | 0.00140±0.00023 |
| S | 0.00024±0.00008 | 0 | 0.00011±0.00006 | 0.00162±0.00062 |
| N | 0.00051±0.00020 | 0 | 0.00019±0.00018 | 0.00407±0.00157 |
| M | / | / | / | / |
| E | / | / | / | / |

Note: \*d., distance;

### Amino acid residues alignment

CDS of polyprotein encodes many proteins, mainly soluble enzymes, which play an important role in the infection cycle of a virus. Its precursor contained the most amino acid residues, up to 7096. There are 22 variations between EPI_ISL_403928 and the other 2019-nCoVs (**Fig 2**).

**Fig 2.**
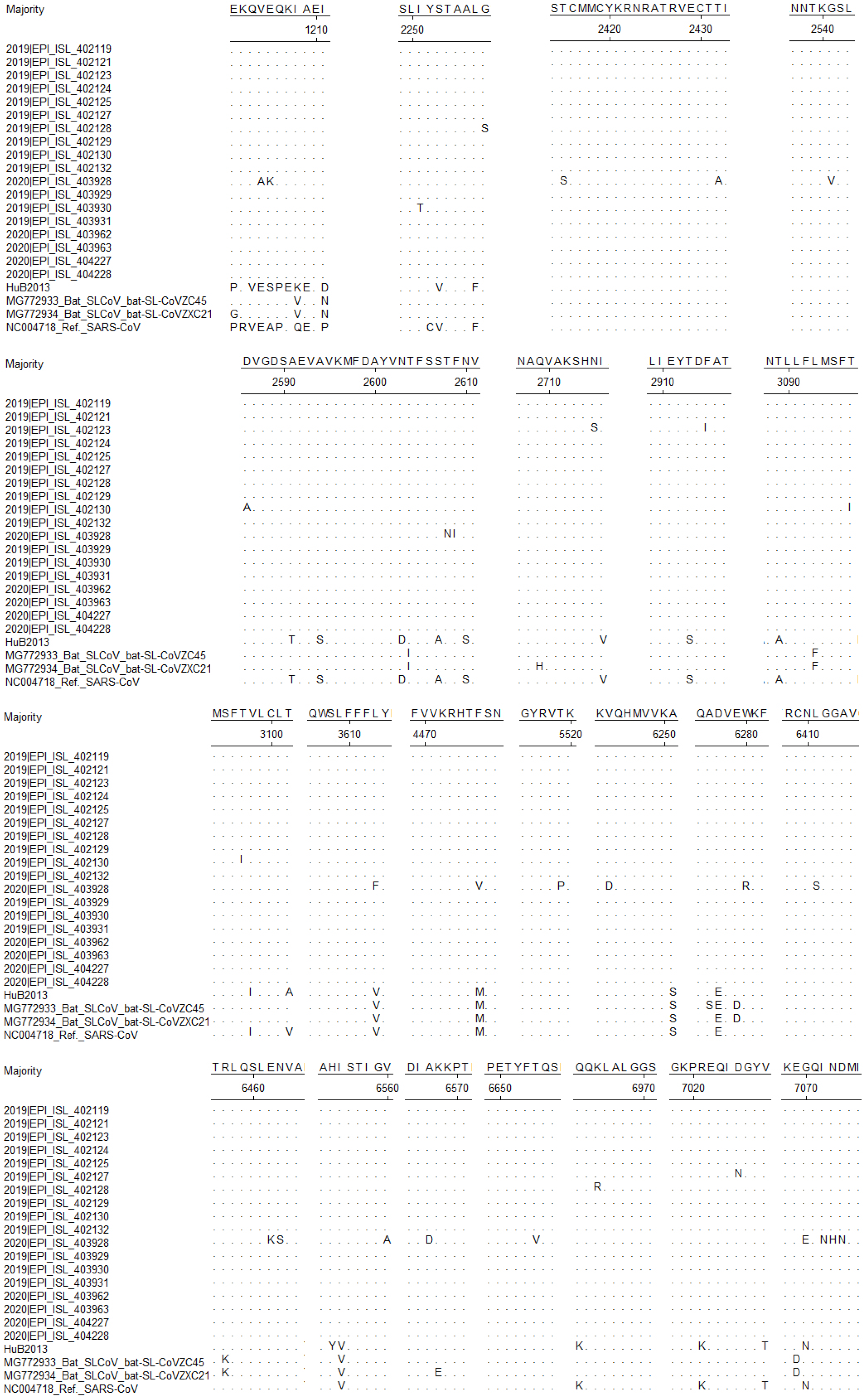
Alignment for amino acid residues of polyprotein of 2019-nCoV. Due to the limited space, only 18 isolates of 2019-nCoV were displayed. Four lines at the bottom were used as references, especially the SARS-CoV (accession NC_004718), to show the correspondence sites of amino acid residues between 2019-nCoV and the known *Betacoronavirus*. Also because of the limited space, the same sites of amino acid residues as EPI_ISL_403928 were omitted.

Two important structural proteins, S and N, are composed of 1273 and 419 amino acid residues, respectively. When compared with the other 2019-nCoVs, EPI_ISL_403928 has four variations in S protein (T572I, G799V, F800C and N801K) and two variations in N protein (A414C and D415I). M and E proteins show no variation so far.

### Evolutionary rate and tMRCA

Nucleotide substitution rates of the whole genome and CDS for different proteins differed from each other (**Tab 2**). The substitution rate from high to low was 1·05 × 10^−2^ (nucleotide substitutions/site/year, with 95% HPD interval being 6.27 × 10^−4^ to 2.72 × 10^−2^, similarly hereinafter) for N, 5.34 × 10^−3^ (5.10 × 10^−4^, 1.28 × 10^−2^) for S, 1.69 × 10^−3^ (3.94 × 10^−4^, 3.60 × 10^−3^) for P, and 1.65 × 10^−3^ (4.47 × 10^−4^, 3.24 × 10^−3^) for whole genome. It is estimated that the most recent common ancestor (tMRCA) of 2019-nCoVs existed about 0.253-0.594 year (somewhat, 91-214 days) ago, or their evolutionary divergence appeared 91-214 days ago. However, M and E demonstrated no divergences up to this point.

**Tab 2.** Evolutionary molecular clock related parameters of 24 isolates of 2019-nCoV.

Gp, group. Gp 1 contains only one genome, EPI\_ISL\_403928, while Gp 2 contains all sequences except EPI\_ISL\_403928. **Tab 2. Evolutionary molecular clock related parameters of 24 isolates of 2019-nCoV**
|  | TMRCA <sup>a</sup> |  |  |  | Nucleotide substitution rate |  |  |  |
| --- | --- | --- | --- | --- | --- | --- | --- | --- |
|  | mean | HPDL <sup>b</sup> | HPDU <sup>c</sup> | ESS <sup>d</sup> | mean | HPDL | HPDU | ESS |
| Genome | 0.594 | 0.198 | 1.152 | 417.2 | 1.65 E-3 | 4.47 E-4 | 3.24 E-3 | 369.2 |
| P | 0.589 | 0.161 | 1.207 | 371.6 | 1.69 E-3 | 3.94 E-4 | 3.60 E-3 | 275.0 |
| S | 0.253 | 6.67 E-2 | 0.599 | 345.9 | 5.34 E-3 | 5.10 E-4 | 1.28 E-2 | 199.7 |
| N | 0.325 | 6.67 E-2 | 0.846 | 611.6 | 1.05 E-2 | 6.27 E-4 | 2.72 E-2 | 447.8 |
| M | / | / | / | / | / | / | / | / |
| E | / | / | / | / | / | / | / | / |

Note: a, the most recent common ancestor;
b, lower of 95% HPD confidence interval;
c, upper of 95% HPD confidence interval;
d, effective sample sizes.

## Discussion

In this study, we analyzed the whole genome and the CDS of several key proteins of 2019-nCoVs. 2019-nCoVs are the pathogens that cause severe pneumonia in human in Wuhan recently. It is found one of the isolates (EPI_ISL_403928) has obvious variations in whole genome and CDS of P, S and N proteins, which result in the substitution of several amino acids.

We also analyzed the nucleotide substitution rate in the whole genome level and the CDS of some proteins during their evolution process. There is no variation in CDS of M and E proteins. CDS of N protein has the highest rate of base substitution (1·05 × 10^−2^ substitutions/site/year) and whole genome has the lowest (1.65 × 10^−3^ substitutions/site/year). The divergence time of the viruses deduced is about 91-214 days before the submission of the first isolate. Their earliest sampling time was 8 December, 2019, and 91-214 days before that day, the viruses infected human or host animals and began to diverge in the evolution process. It is not clear if the viruses began to mutate before or after infecting human beings. There is also a possibility there were two strains evolved coincidentally, one for human and one for host animals that evolved more adaptive to human and then infected human. Our analysis indicates that the outbreak occurred a few months earlier than it was officially reported, and there might be recessive infection and underdetection in the population. Therefore, the seafood market in Wuhan, which is believed to be the first spot of the outbreak, may or may not be the case.

Human recessive infection of *Betacoronavirus* does exist. For example, it occurs in the population for HKU1, another member of β-CoVs. HKU1 is related to a variety of children’s and adult diseases and distributed globally [16]. In the United States, 0.5% HKU1 positive rate was detected in 15 287 respiratory samples during 2009-2013 [17]; in Kenya, Africa, 2.1% of the 417 respiratory samples collected in 2009-2012 were infected by HKU1 [18]; in Southeast Asia, the positive rate of HKU1 was 1.1% in 2060 adult acute respiratory samples from Malaysia in 2012-2013 [19]. The positive detection rate was 1.9% in Japan and 4342 from 2010 to 2013, and 2.5% in Korea [20,21]. Recessive HKU1 infection was common and many patients with HKU1 infection were found in routine examinations [24–25]. Cases of recessive 2019-nCoV infection have also been reported in succession. Recessive infection is one of the reasons for underdetection.

All human CoVs (HCoVs) is mainly of zoonotic origin, and most likely originate from bats [9]. The common scenario of CoV evolution then involves past transitions into intermediate hosts such as livestock, which have closer interaction with human and may carry a diversity of viruses including variants directly related to ancestral strains [26,27]. For this outbreak, there is evidence for specific wild animals being intermediate hosts in the seafood market in Wuhan, similar to the outbreak of SARS-CoV in Guangdong Province of China in 2002-2003 [28]. People engaged in hunting and management of such wild animals are at high risk of infection, likely live in mountain or rural areas and are more likely to be undetected when having such an infection for various reasons.

## Conclusions

Our study suggests that there are at least two different viral strains of 2019-nCoV infecting human and human infection occurred a few months earlier than the outbreak being officially announced. Recessive infection and underdetection can cause a delay in response. The seafood market in Wuhan may or may not be the first spot of this outbreak. A large number of viruses carried by wild animals bring many uncertainties to the emerging infectious diseases (EIDs). In order to effectively control these EIDs, it is necessary to strengthen interdisciplinary cooperation and communication among human, animal and environmental health investigators based on the One Health concept, so as to detect and identify pathogens as early as possible, find patients and report epidemics, and effectively control the spread of EIDs through timely isolation and prevention measures and observation of close contacts.

## List of abbreviations

CoVs: Coronaviruses
2019-nCoV: 2019-novel coronavirus
SARS-CoV: severe acute respiratory syndrome coronavirus
MERS-CoV: Middle East respiratory syndrome coronavirus
CDS: coding sequence
tMRCA: the most recent common ancestor
GISAID: the Global Initiative on Sharing Avian Influenza Database
ESSs: Effective sample sizes

## Declarations

### ETHICS APPROVAL AND CONSENT TO PARTICIPATE

This study is a serial of phylogenetic analyses based on large scale of existing gene sequences; all these sequences can be searched and downloaded from two public databases, the NCBI Influenza Virus Sequence Database and the Global Initiative on Sharing Avian Influenza Data (GISAID) database. No institutional review board approval was required from the research ethics committee of School of Public Health, Fudan University, and animals’ ethics approval was applicable neither.

### CONSENT FOR PUBLICATION

All authors have approved publishing this paper in *Infectious Diseases of Poverty*, and there are no patients involved in this study.

### AVAILABILITY OF DATA AND MATERIALS

Not applicable.

### AUTHORS’ CONTRIBUTIONS

All authors made significant contributions to the conception, data acquisition, analysis and drafting of this manuscript and approve the final version submitted. C. X. and Q. J. conceived and designed the project. Y. C. and Q. J. developed the research question. C. X. and L. J. collected the sequences and calculated them. All members of the group contributed to the analysis design and interpretation of the data.

## ACKNOWLEDGEMENTS

We acknowledge the contributions of scientists and researchers from all over the world for depositing the genomic sequences of influenza viruses in the Global Initiative on Sharing All Influenza Data (GISAID) EpiFlu- and the nucleotide database of the National Center for Biotechnology Information (NCBI). We acknowledge these two databases for permitting us to use these genomic sequences freely and conveniently.

